# *Paso doble*: A two-step Late Pleistocene range expansion in the Tyrrhenian tree frog *Hyla sarda*

**DOI:** 10.1101/2020.07.21.213363

**Authors:** Giada Spadavecchia, Andrea Chiocchio, Roberta Bisconti, Daniele Canestrelli

**Author notes:** These authors contributed equally to this work (Roberta Bisconti, Daniele Canestrelli). **Correspondence:** Roberta Bisconti.

## Abstract

The Tyrrhenian tree frog, *Hyla sarda*, is an amphibian endemic to the Tyrrhenian islands (Western Mediterranean). Previous investigations of its Pleistocene evolutionary history suggested that it colonised the northern portion of its current range, through a spatial diffusion process from the Sardinia island, during the last glaciation. However, southern and northern portions of the species’ range experienced markedly different climatic conditions during the Late Pleistocene, suggesting the possibility of an unusual two-step process of demographic expansion. Here, we use Bayesian phylogeographic approaches to locate the ancestral area in Sardinia and to characterise better the demographic component of this expansion event. These analyses located the ancestral area for *H. sarda* populations along the central-eastern coast of the Sardinia island, within an area previously shown to host suitable bioclimatic conditions for *H. sarda* populations throughout the Late Pleistocene. Historical demographic reconstructions clearly showed that a two-step process of demographic growth fits well the data, with northern populations expanding later than Sardinia populations. The harsher climatic conditions occurred in northern islands during the glacial epoch, as compared to Sardinia, likely delayed tree frog colonisation of northern territories, and the associated demographic growth.

## Introduction

Species showing unusual phylogeographic patterns might provide unique opportunities to explore the role of a wide range of ecological and evolutionary processes in shaping spatial patterns of biological diversity. However, to fully exploit these opportunities, we need a thorough understanding of the underlying demographic histories (e.g. Barbosa et al. 2017; Canestrelli et al. 2010). During the past four decades, phylogeographic investigations of temperate species in the Western Palearctic have documented widespread demographic and range contractions during glacial epochs, followed by expansions during subsequent interglacials (Hewitt 2000; 2004a; 2004b; 2011a; Taberlet et al. 1998). Although shared among a wide range of organisms, from all sub-regions of the Western Palearctic and beyond (Hewitt 2011b), this general scenario is not devoid of exceptions. One of these remarkable exceptions is represented by instances of glacial demographic and range expansion for temperate species, a reverse expansion-contraction scenario initially proposed for the Tyrrhenian tree frog, *Hyla sarda* (Bisconti et al. 2011a; 2011b), and recently used to explain phylogeographic patterns in other organisms (e.g. Porretta et al. 2012; Senczuk et al. 2019).

The Tyrrhenian tree frog is a small, cryptically coloured amphibian, endemic to the Western Mediterranean islands of Sardinia and Corsica, and the Tuscan archipelago. It is a temperate species, widespread from the sea level up to 1200 meters a.s.l., although it is markedly more abundant along the coastal areas (Lanza et al. 2007). *H. sarda* has primarily aquatic habits, living close to lentic freshwater habitats, such as pools and temporary ponds. Previous studies of its Pleistocene evolutionary history (Bisconti et al. 2011a; 2011b) showed that, contrary to what has been found in most temperate species studied to date, including amphibians (Zeisset et al. 2008), this tree frog likely initiated a phase of major demographic expansion in the middle of the last glaciation. This event, likely promoted by a glaciation-induced increase in lowland areas during the marine regression, also allowed *H. sarda* to colonise the northern island of Corsica and the Tuscan archipelago, from an ancestral area in Sardinia, taking advantage of a wide and persistent land bridge connecting the Sardinia to Corsica throughout the glacial epoch (Bisconti et al. 2011a; 2011b).

Two main points left open by previous studies were: i) the geographic location of the ancestral area in northern Sardinia, and ii) the timing of the northward range expansion into Corsica and the Tuscan archipelago. In fact, quantitative analyses were not conducted to attempt locating the ancestral area. Most importantly, while the historical demographic reconstruction clearly showed a full-glacial development of the expansion event (Bisconti et al. 2011a), paleoclimatic reconstructions for the Western Mediterranean indicated substantially harsher climatic conditions in central and northern Corsica than in Sardinia, during the last glaciation (Kuhlemann et al. 2008; Hughes &Woodward 2017; Pascucci et al. 2014). In turn, this paleoclimatic reconstruction for the last glaciation suggests an intriguing scenario, whereby tree frog population expansion in Corsica and the Tuscan archipelago might have been delayed, as compared to Sardinia populations, leading to a two-step expansion process. A first step, promoted by the glaciation-induced widening of coastal areas in Sardinia (as suggested by Bisconti et al. 2011a), would have been followed by a second step leading tree frog populations to complete the northward colonisation, following post-glacial climatic amelioration in northern areas.

Here, we explore this (revised) phylogeographic scenario, through a Bayesian phylogeographic analysis, coupled with new historical demographic assessments. We carried out the demographic component of the analysis separately for the source (northern Sardinia) and the recolonised (Corsica and the Tuscan archipelago) populations, with the following rationale. Although they belong to the same phylogeographic lineage (Bisconti et al. 2011a), their demographic history might have been differently shaped by the recent paleoclimatic changes of the respective geographic ranges. If this was the case, their demographic reconstructions would differ accordingly. Instead, in case they actually behaved as a single demographic unit in response to such changes (following Bisconti et al. 2011a), the respective demographic trends would be broadly similar, as they would represent two independent samples of the same demographic unit.

## Materials and methods

We collected tissue samples from 81 individuals of *H. sarda*. Tree frogs were sampled by toe-clipping after anaesthetization in a 0.1% solution of MS222 (3-aminobenzoic acid ethyl ester) and then released at the respective collection site, while tissue samples were stored in 95% alcohol. Sampling and experimental procedures were approved by the Italian Ministry of Environment ‘MATTM’ (protocol #8275), and Prefecture of Corsica (#2A20180206002 and #2B20180206001). Samples collected for the present study were complemented with data from northern Sardinia, Corsica and the Tuscan archipelago, collected by previous studies (Bisconti et al. 2011a), allowing an overall sample size of 171 individuals from 18 sampling locations (Table 1, Figure 1A, and Table S1). Instead, data from central and southern Sardinia from the previous studies were not considered here, as these areas were shown to be populated by distinct mitochondrial sub-lineages (Bisconti et al. 2011a).

**Table 1.** Sampling location and sample size (n) for the 18 populations of *Hyla sarda* analysed for the present study.

| Island |  | Locations | n | Latitude N | Longitude E |
| --- | --- | --- | --- | --- | --- |
| Sardinia | 1 | Monteferro | 14 | 40°12' | 08°35' |
|  | 2 | Asinara | 2 | 41°03' | 08°14' |
|  | 3 | Cala Ginepro | 8 | 40°26' | 09°47' |
|  | 4 | Siniscola | 25 | 40°34' | 09°46' |
|  | 5 | Posada | 6 | 40°38' | 09°45' |
|  | 6 | Vallicciola | 5 | 40°51' | 09°09' |
|  | 7 | Luogosanto | 9 | 41°03' | 09°12' |
|  | 8 | Stazzo<br>Pulcheddu | 6 | 41°09' | 09°21' |
|  | 9 | Porto Pollo | 9 | 41°11' | 09°19' |
| Corsica | 10 | Etang de<br>Canettu | 6 | 41°25' | 09°13' |
|  | 11 | T10 | 10 | 41°26' | 09°12' |
|  | 12 | L'Ospedale | 12 | 41°39' | 09°11 ' |
|  | 13 | Aleria | 11 | 42°06' | 09°31' |
|  | 14 | Curzo | 9 | 42°19' | 08°40' |
|  | 15 | San Giuliano | 7 | 42°16' | 09°31' |
|  | 16 | Biguglia | 3 | 42°38' | 09°26' |
| Capraia | 17 | Capraia | 5 | 43°02' | 09°50' |
| Elba | 18 | Elba | 25 | 42°47' | 10°15' |

**Figure 1.**
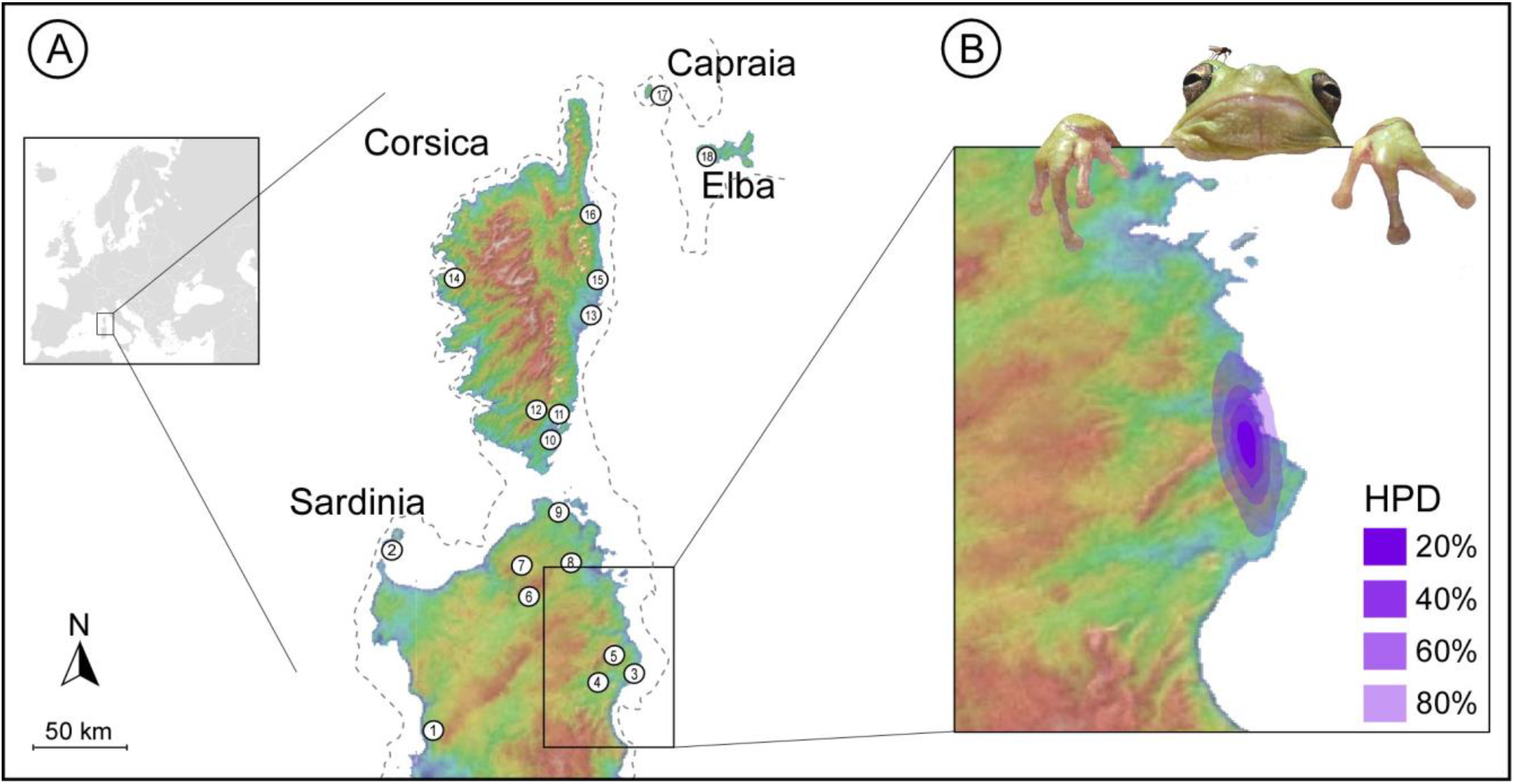
A) Study area and geographic location of the 18 sampled populations of *H. sarda*. The dashed line shows the approximate location of the coastline during the last glacial maximum (Thiede, 1978). B) Highest posterior density (HPD) regions of the ancestral area of *H. sarda*, based on a Bayesian phylogeographical analysis carried out in Beast.

Following a step of tissue fragmentation and digestion with proteinase K, total DNA was extracted using the standard phenol-chloroform method (Sambrook et al. 1989). Two mitochondrial gene fragments were amplified by polymerase chain reactions (PCRs): cytochrome b (cytb) and NADH dehydrogenase subunit 1 (ND1). Primers used and PCR cycling conditions were the same described in Bisconti et al. (2011a). All sequencing procedures were performed by Macrogen Inc. (www.macrogen.com). The multi-purpose sequence analysis suite GeneStudio (available at www.genestudio.com) was used to check the electropherograms by eye, and to generate multiple sequence alignments. All sequences obtained were deposited in the GenBank database (Table S1). For all downstream analyses, the two gene fragments were concatenated using DnaSp 6.11.01 (Rozas et al. 2017). The models of nucleotide substitution that best fit the data (HKY for both CytB and ND1) were identified by means of PartitionFinder 2.1.1 (Lanfear et al. 2016) using the Bayesian information criterion (Schwarz 1987).

The geographic location of the ancestral area for the inferred northward expansion process (Bisconti et al. 2011a), was investigated by estimating a Bayesian phylogeographic diffusion model in continuous space (BP), as implemented in Beast 1.10.4 (Suchard et al. 2018). The analysis was run using clock models and substitution models unlinked across all partitions. A Bayesian skyline was used as coalescent prior, and the strict molecular clock was enforced, as it is generally a good approximation for analyses at the intrapopulation level (Yang 2006) and because, by simplifying the coalescent model, it helps analyses to converge (Heled 2010). The substitution rate was set to 1.37 × 10^−8^, as estimated for *H. sarda* by the previous study (Bisconti et al. 2011a). After some exploratory runs, the final analyses were run for 100 million generations, sampled every 1000^th^ generation, using a relaxed random walk diffusion model (RRWs) with Cauchy distribution (Lemey et al. 2009). Convergence among runs, effective sample size (ESS) values, and the appropriate burn-in were evaluated using the software Tracer 1.7.1 (Rambaut et al. 2018). Finally, we visualised the ancestral area, and its changes through time, using the Time Slicer function implemented in Spread 1.0.7 (Bielejec et al. 2011), which estimate Highest Posterior Density (HPD) regions for the parameter of interest, based on the full tree forest.

Historical demographic trends were estimated through the Bayesian skyline plot (BSP) model implemented in Beast. These analyses were performed using the same settings as above, ten piecewise constant intervals, and a uniform prior distribution for the population size parameter. Since preliminary BSP analyses suggested demographic growth following a phase of constant population size, for both Sardinia and Corsica populations, we estimated the transition time between these two demographic ‘epochs’ (i.e. an epoch of constant population size followed by an epoch of population growth) by means of a Two-Epoch analysis in Beast (Crandall et al. 2012). For this analysis, custom *.xml* files were prepared following suggestions provided by Crandall et al. (2012). Preliminary analyses were run using both an exponential and a logistic model population growth. Estimates of the transition time did not differ appreciably with the two models. However, the logistic model yielded comparatively inferior performance statistics and is not reported here (available upon request). These analyses were run for 100 million generations, sampled every 1000^th^ generation. All the analyses with Beast were run twice and then combined to generate the final results, using LogCombiner 1.10.4.

## Results

We obtained concatenated sequences 1226 bp long for all the individuals analysed, leading to a final alignment including 171 individuals (81 from this study, 90 from Bisconti et al. 2011a). Sequences represented uninterrupted open reading frames, with no gaps or premature stop codons, indicating they are functional mitochondrial DNA copies. All the analyses carried out with BEAST converged to a stationary distribution, with high effective sample size values (>200) for all the parameters of interest.

The Bayesian phylogeographic analysis identified the ancestral area of *H. sarda* populations along a small stretch of the middle eastern coastal region in Sardinia (Figure 1B). According to this analysis, the species remained in this area until the last glacial epoch, when it started spreading in Sardinia and then to the north, toward the Corsica island and the Tuscan archipelago (see Video S1).

The historical demographic reconstructions carried out by means of Bayesian skyline plot analyses are shown in Figure 2A. Both for populations in Sardinia and for populations in Corsica and the Tuscan Archipelago, these analyses identified an initial phase of demographic stasis, followed by marked population growth. However, the timeline of the inferred demographic trends was markedly different. In Sardinia, the time of the most recent common ancestor (TMRCA) was dated at 129 thousand years ago (kya; 95%HPD: 71 – 216 kya), whereas the transition time between the two demographic epochs (i.e. constant *vs* growth; see Figure 2B) was estimated to have occurred 58 kya (95%HPD: 40 – 79 kya). Instead, in Corsica and the Tuscan Archipelago, the TMRCA was estimated at 59 kya (95%HPD: 24 – 119 kya), while the transition time to the expansion epoch was dated at 34 kya (95%HPD: 3 – 63 kya).

**Figure 2.**
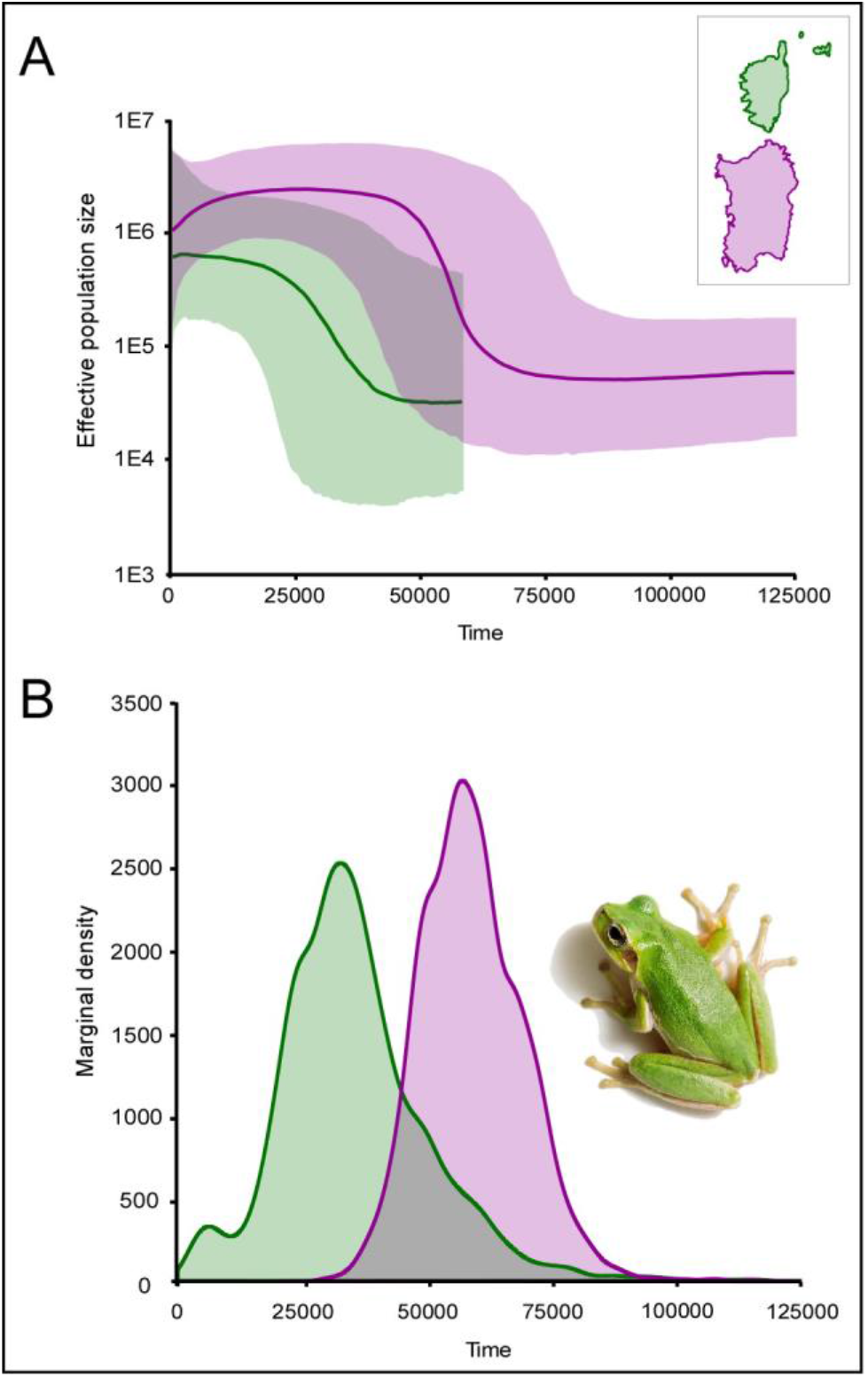
Historical demographic reconstructions for the Sardinian populations (purple), and the Corsica and Tuscan archipelago populations (green). A) Bayesian skyline plots showing the effective population size change through time. Median estimates and 95% highest posterior density regions are shown (continuous line and shaded areas, respectively). B) Marginal posterior probability distributions for the transition time to exponential growth from the two-epoch models.

## Discussions

The previous phylogeographic investigation suggested that glaciation-induced widening of coastal plains might have promoted a demographic and range expansion of the Tyrrhenian tree frog, during the last glaciation (Bisconti et al. 2011a). On the other hand, fine-scale paleoenvironmental reconstructions for the Late Pleistocene of the Tyrrhenian islands, support this scenario for the Sardinia island, but not for Corsica and the other northern islands. Here, the glacial climate was harsher and coastal widening was limited, compared to Sardinia (Thiede 1978; Kuhlemann et al. 2005 and 2008; Forzoni et al. 2015). Our Bayesian analyses of the recent evolutionary history of *H. sarda* populations allowed us to reconcile these phylogeographic and paleoenvironmental perspectives and suggests an intriguing new scenario for the persistence of temperate species in the Mediterranean region, over glacial-interglacial cycles.

Our results support previous inference suggesting that *H. sarda* (re) colonised its entire range during a Late Pleistocene expansion. However, they also suggest that this major expansion phase did not occur as a single demographic event, but most probably through two sequential – albeit tightly linked - expansion steps. The first step allowed *H. sarda* to (re) colonise the rest of the Sardinia island during the last glaciation, and to expand into the extensive coastal plain opened by the glaciation induced sea-level drop north of this island. The second step, which most probably initiated close in time to the end of the last glacial maximum, allowed the species to expand northward from northern Sardinia, and to colonise the entire island of Corsica and the Tuscan archipelago.

The source area of the Late Pleistocene expansion of *H. sarda* populations was identified in a narrow coastal region in north-east Sardinia (Figure 1B). During the last glaciation, the widening of the coastal lowland adjoining this area was conspicuous (see Figure 1B), and it progressively gained a direct connection with the vast lowland which gradually connected northern Sardinia with southern Corsica. Likewise, previously estimated species distribution models (Bisconti et al. 2011a) indicated high bioclimatic suitability of this area for *H. sarda*, under both current and glacial bioclimatic scenarios. Hence, phylogeographic and paleoclimatic data converge in identifying this area as a suitable candidate ancestral area, and source area for the subsequent demographic expansion.

In line with previous historical demographic reconstructions carried out at the level of the entire species’ range (Bisconti et al. 2011a), our Bayesian skyline plot analysis of Sardinian populations suggested that the expansion event initiated in the middle of the Late Pleistocene (58 kya). To some extent, this time estimate might be an overestimate, owing to the time-dependency of the molecular clock (Ho et al. 2015), and the application of a molecular clock rate calibrated using the Messinian salinity crisis (i.e. 5.3 million years ago). Yet this time estimate matches considerably well with a phase of major sea-level drop (occurring about 65 kya; Spratt & Lisiecki, 2016), and a consequent widening of coastal habitats. Most importantly, however, the estimated region of 95% highest posterior probability density for this event (40 – 79 kya), rule out the possibility of a post-glacial, or even a late-glacial, initiation of the demographic expansion in Sardinia. Thus, phylogeographic and paleoclimatic data converge in identifying the middle of the Late Pleistocene as the most likely time frame for the expansion event in Sardinia.

In Corsica and the Tuscan archipelago, however, the expansion event occurred later. For these populations, the inferred TMRCA is significantly more recent than for Sardinian populations, and closely matches the transition time between the two demographic epochs in Sardinia (median estimates: 59 *vs* 58 kya, respectively). Importantly, although surrounded by a non-negligible uncertainty (95%HPD: 24 – 119 kya), this event appears unlikely to have followed the last glacial maximum (LGM; ~23 kya; Kuhlemann et al. 2008). Instead, the same cannot be said for the expansion event. Since the TMRCA, a phase of demographic stability lasting about 25 kya preceded the demographic expansion. Although the median estimate for the transition time between these two epochs (34 kya) precedes the LGM, the 95% highest posterior density region for this event (3 – 63 kya; see also Figure 2B) largely incorporates the LGM and subsequent post-glacial times. Moreover, since this is the most recent event inferred from our data, we can expect it to be most affected by the time-dependency of the molecular clock, and the consequent inflation of recent time estimates (see Membrebe et al. 2019 for an estimate of the effect size in a case where both root calibration and dated tips were available). Lastly, when also considering the particularly harsh climate at the LGM in Corsica, due to polar air incursions (Kuhlemann et al. 2008), and the abrupt transition to post-glacial environmental conditions in this area (Kuhlemann et al. 2005 and 2008; Forzoni et al. 2015), a scenario considering a peri-glacial initiation and post-glacial development of the range expansion in Corsica appears the most plausible for a thermophilic species, as is *H. sarda*.

## Conclusions

Our study allowed to improve previous estimates of the evolutionary history of the Tyrrhenian tree frog *H. sarda*, by 1) locating the source area of its major range expansion in the Late Pleistocene, and 2) identifying a previously undetected component of the associated demographic trends. On the one hand, the inferred two-step expansion model is unprecedented for temperate species in the Mediterranean region and will deserve further evaluation in other temperate species. These studies should be focused on species both from insular geographic settings and from mainland areas close to coastal regions which have been particularly affected by glacial widenings of lowland habitats (e.g. the Adriatic Sea, central Mediterranean). On the other hand, by providing unusual resolution to the temporal and geographic components of its recent evolutionary history, our results place *H. sarda* in an excellent position to become a prominent case for the study of the genomic and phenotypic legacy of Late Pleistocene range dynamics of island species.

Finally, although previous studies showed no significant discrepancy between nuclear and mtDNA data (Bisconti et al. 2011a), results from the present study should be confirmed with in-depth analyses of the nuclear genome. Whole-genome sequencing will be useful to precisely calibrate the molecular clock, to confirm the demographic dynamics that emerged in the present study, and to investigate the evolutionary implications of these dynamics.

## Acknowledgements

We are grateful to Alessandro Carlini, Giacomo Grignani, Lorenzo Latini, Anita Liparoto, Armando Macali, for assistance with sampling and the experimental procedures. The research was supported by a grant from the Italian Ministry of Education, University and Research (PRIN project 2017KLZ3MA).

## Author contributions

R.B. and D.C. designed the study; G.S., A.C., R.B. and D.C. performed research; G.S. and D.C. analyzed data; G.S., R.B. and D.C. wrote the paper; G.S., A.C., R.B. and D.C. discussed and approved the final version of the manuscript.

## Conflict of Interest

The authors declare that the research was conducted in the absence of any commercial or financial relationships that could be construed as a potential conflict of interest.

## SUPPORTING INFORMATION

Additional Supporting Information may be found in the online version of this article at the publisher’s web-site:

**Table S1.** Complete dataset generated and used in this study.

**Video S1.** Animation of the spatial diffusion through time of *Hyla sarda* in virtual globe software (GoogleEarth).

## Notes

### Competing Interest Statement

The authors have declared no competing interest.

## References

Barbosa S, Paupério J, Herman JS, Ferreira CM, Pita R, Vale-Gonçalves HM, & Mira A. 2017. Endemic species may have complex histories: Within-refugium phylogeography of an endangered Iberian vole. Molecular ecology, 26(3), 951–967.

Bielejec F, Rambaut A, Suchard MA, Lemey P. 2011. SPREAD: spatial phylogenetic reconstruction of evolutionary dynamics. Bioinformatics, 27(20), 2910–2912.

Bisconti R, Canestrelli D, Colangelo P, Nascetti G. 2011a. Multiple lines of evidence for demographic and range expansion of a temperate species (*Hyla sarda*) during the last glaciation. Molecular Ecology, 20(24), 5313–5327.

Bisconti R, Canestrelli D, Nascetti G. 2011b. Genetic diversity and evolutionary history of the Tyrrhenian treefrog Hyla sarda (Anura: Hylidae): adding pieces to the puzzle of Corsica–Sardinia biota. Biological Journal of the Linnean Society, 103(1), 159–167.

Canestrelli D, Aloise G, Cecchetti S, Nascetti G. 2010. Birth of a hotspot of intraspecific genetic diversity: notes from the underground. Molecular Ecology, 19(24), 5432–5451.

Crandall ED, Sbrocco EJ, DeBoer TS, Barber PH, Carpenter KE. 2012. Expansion dating: calibrating molecular clocks in marine species from expansions onto the Sunda Shelf following the Last Glacial Maximum. Molecular Biology and Evolution, 29(2), 707–719.

Forzoni A, Storms JEA, Reimann T, Moreau J, Jouet G. 2015. Non-linear response of the Golo River system, Corsica, France, to Late Quaternary climatic and sea level variations. Quaternary Science Reviews, 121, 11–27.

Heled J. 2010. Extended Bayesian skyline plot tutorial. Available from http://beast-mcmc.googlecode.com

Hewitt G. 2000. The genetic legacy of the Quaternary ice ages. Nature, 405(6789), 907.

Hewitt G. 2004a. Genetic consequences of climatic oscillations in the Quaternary. Philosophical Transactions of the Royal Society of London. Series B: Biological Sciences, 359(1442), 183–195.

Hewitt G. 2004b. The structure of biodiversity–insights from molecular phylogeography. Frontiers in zoology, 1(1), 4.

Hewitt G. 2011a. Mediterranean Peninsulas—the evolution of hotspots. In: Biodiversity Hotspots (eds Zachos FE and Habel JC), pp. 123–148. Springer, Amsterdam.

Hewitt G. 2011b. Quaternary phylogeography: the roots to hybrid zones. Genetica, 139, 617–638.

Ho SY, Duchêne S, Molak M, Shapiro B. 2015. Time-dependent estimates of molecular evolutionary rates: evidence and causes. Mol Ecol. 2424:6007 – 6012. doi: 10.1111/mec.13450

Hughes PD, Woodward JC. 2017. Quaternary Glaciation in the Mediterranean Mountains. Geological Society, London, Special Publications, 433, 1–23.

Kuhlemann J, Frisch W, Székely B, Dunkl I, Danišik M, Krumrei I. 2005. Würmian maximum glaciation in Corsica. Austrian Journal of Earth Sciences, 97, 68–81.

Kuhlemann J, Rohling EJ, Krumrei I, Kubik P, Ivy-Ochs S, Kucera M. 2008. Regional synthesis of Mediterranean atmospheric circulation during the last glacial maximum. Science, 321, 1338–1340.

Lanza B, Andreone F, Bologna MA, Corti C, Razzetti E. 2007. Fauna d’Italia Amphibia. Calderini, Bologna.

Lanfear R, Frandsen PB, Wright AM, Senfeld T, Calcott B. 2016. PartitionFinder 2: new methods for selecting partitioned models of evolution for molecular and morphological phylogenetic analyses. Molecular biology and evolution.

Lemey P, Rambaut A, Drummond AJ, Suchard MA. 2009. Bayesian phylogeography finds its roots. PLoS computational biology, 5(9), e1000520.

Membrebe JV, Suchard MA, Rambaut A, Baele G, Lemey P. 2019. Bayesian inference of evolutionary histories under time-dependent substitution rates. Molecular biology and evolution, 36(8), 1793–1803.

Pascucci V, Sechi D, Andreucci S. 2014. Middle pleistocene to holocene coastal evolution of NW sardinia (Mediterranean Sea, Italy). Quaternary International, 328, 3–20.

Porretta D, Mastrantonio V, Bellini R, Somboon P, Urbanelli S. 2012. Glacial history of a modern invader: phylogeography and species distribution modelling of the Asian tiger mosquito Aedes albopictus. PloS one, 7(9).

Rambaut A, Drummond AJ, Xie D, Baele G, Suchard MA. 2018. Posterior summarization in Bayesian phylogenetics using Tracer 1.7. Systematic biology, 67(5), 901–904.

Rozas J, Ferrer-Mata A, Sánchez-DelBarrio JC, Guirao-Rico S, Librado P, Ramos-Onsins SE, Sánchez-Gracia A. 2017. DnaSP 6: DNA sequence polymorphism analysis of large data sets. Molecular Biology and Evolution, 34(12), 3299–3302.

Sambrook J, Fritsch EF, Maniatis T. 1989. Molecular cloning: a laboratory manual (Ed. 2). Cold spring harbor laboratory press.

Schwarz G. 1987. Estimating the dimension of a model. The annals of statistics, 6(2), 461–464.

Senczuk G, Harris DJ, Castiglia R, Litsi Mizan V, Colangelo P, Canestrelli D, Salvi D. 2019. Evolutionary and demographic correlates of Pleistocene coastline changes in the Sicilian wall lizard Podarcis wagleriana. Journal of Biogeography, 46(1), 224–237.

Spratt RM, Lisiecki LE. 2016. A Late Pleistocene sea level stack. Climate of the Past, 12(4), 1079–1092.

Suchard MA, Lemey P, Baele G, Ayres DL, Drummond AJ, Rambaut A. 2018. Bayesian phylogenetic and phylodynamic data integration using BEAST 1.10. Virus Evolution, 4(1), vey016.

Taberlet P, Fumagalli L, Wust-Saucy AG, Cosson JF. 1998. Comparative phylogeography and postglacial colonization routes in Europe. Molecular ecology, 7(4), 453–464.

Thiede J. 1978. A glacial Mediterranean. Nature, 276, 680–683.

Yang Z: Computational Molecular Evolution. Oxford University Press 2006, Oxford, UK

Zeisset I, Beebee TJC. 2008. Amphibian phylogeography: a model for understanding historical aspects of species distributions. Heredity, 101(2), 109–119.

